# Spider mites collectively avoid plants with cadmium irrespective of their frequency or the presence of competitors

**DOI:** 10.1101/2023.08.17.553707

**Authors:** Diogo Prino Godinho, Inês Fragata, Maud Charlery de la Masselière, Sara Magalhães

## Abstract

Accumulation of heavy metals by plants can serve as a defence against herbivory. Herbivores, in turn, may avoid feeding on contaminated tissues. Such avoidance, however, may hinge upon the specific conditions faced by herbivores. Here, we tested whether the spider mite *Tetranychus urticae* avoids tomato plants contaminated with cadmium in presence of conspecifics or heterospecifics and depending on the frequency of contaminated plants. We show that individual spider mite females do not preferentially move to leaf tissues with or without cadmium, despite clear costs on their performance. However, in a set-up where 200 mites were simultaneously given the choice between four plants with or without cadmium, they collectively avoided plants with cadmium, irrespective of the proportion of plants with cadmium. In addition, *T. urticae* did not discriminate between plants infested with its competitor *T. evansi* and other uncontaminated plants but they preferred plants with competitors when the other plants contained cadmium. Our results show that aggregation may facilitate avoidance of contaminated plants. They also indicate that cadmium accumulation in plants is a stronger selective pressure than interspecific competition with *T. evansi.* Therefore, collective avoidance of metal-accumulating plants by herbivores is robust to environmental conditions and may have important consequences for species distribution and interactions in metal contaminated sites.

## Introduction

Soils often contain heavy metals, either due to geochemical properties or to human activity (Alloway, 2013; Hou et al., 2017). Some plants, known as hyperaccumulators, can store metals translocated from the soil in their upper parts, up to levels that are toxic to most organisms (Baker, 1987). Such hyperaccumulation often results in a significant reduction in the performance of herbivores that feed on those plants (Boyd & Moar, 1999; Freeman et al., 2007; Kazemi-Dinan et al., 2014; Mohiley et al., 2020; Quinn et al., 2010). This observation is at the basis of the elemental defence hypothesis, which posits that plants use such hyperaccumulated metals as a defence against herbivores (Martens & Boyd, 1994).

Because metal accumulation is detrimental to most herbivores, one would expect the latter to evolve means to overcome this effect. Indeed, some herbivores cope with elemental defences by evolving tolerance mechanisms. Evidence for such adaptation is, however, still scarce (Freeman et al., 2006; Xu et al., 2020), and not universal (Godinho, Branquinho, et al., 2023). Alternatively (or in addition), a common strategy used by herbivores is the avoidance of plants that have accumulated metals (Kazemi-Dinan et al., 2014; Mogren and Trumble, 2010; Vickerman et al., 2003). For example, when given the choice between *Thlaspi caerulascens* plants with high or low zinc concentrations, the desert locust, *Schistocerca gregaria* preferred the latter, a preference pattern that was recapitulated on artificial food supplemented with zinc (Behmer et al., 2005). Another study showed that cadmium modifies the volatiles of *Populus yunnanensis*, and herbivores (the sawfly *Stauronematus compressicornis* and the leaf beetle *Plagiodera versicolora*) use these cues to avoid those plants (Lin et al., 2022).

Herbivores that have been exposed to contaminated food may use this information to ameliorate their choices. Indeed, grass miners (*Chromatomyia milii*) that have fed on grasses (*Holcus lanatus*) with cadmium avoid these plants more than naïve individuals (Pollard & Baker, 1997). However, feeding on metal-contaminated plants may also impair the behavioural decisions of herbivores (Mogren & Trumble, 2010). This, in turn, may affect their ability to avoid contaminated food. Indeed, the ability of flies (*Drosophila melanogaster*) to avoid substrates with lead vanishes when they develop on lead-contaminated food (Peterson et al., 2020). In line with this, another discrimination between plants with or without metals may also be affected by their frequency in the environment. However, to your knowledge, this has not been tested. Apart from affecting the choice between contaminated and non-contaminated food, exposure to heavy metals may also affect other relevant behavioural decisions in herbivores (Oliveira et al., 2022). For example, in presence of competitors, pyramid ants that had ingested selenium had impaired foraging behaviour (De La Riva & Trumble, 2016). Similarly, exposure to zinc compromised the foraging behaviour and predator avoidance in damselfly larvae (Janssens et al., 2014). Because herbivores are often simultaneously exposed to abiotic and biotic pressures, it is key to understand how their interactive effect on herbivore decisions affect their distribution (Tran et al., 2019).

The presence of conspecifics may affect the distribution of herbivores by means of aggregative behaviour (Hunter, 2000; McLellan & Montgomery, 2023). Herbivores may benefit from being with conspecifics, for example to better overcome plant defences (Wertheim et al., 2005). Moreover, behavioural decisions may be affected by the presence of others (Prokopy & Roitberg, 2001). For example, fruit flies (*Bactrocera tryoni*) avoid low-quality food patches more efficiently when in groups (Morimoto et al., 2018). Knowledge that metals affect collective behaviour has only been gathered in zebrafish (Shelton et al., 2023). Whether organisms collectively avoid metal-contaminated food is expected to hinge upon the relative role of acquisition of information and behavioural impairment that such environments represent.

In this study, we test whether herbivorous spider mites avoid plants with cadmium and whether such avoidance hinges on the presence of conspecifics or heterospecifics, or on the frequency of hyperaccumulating hosts. Cadmium is a toxic, non-essential metal widespread in the environment naturally and as an anthropogenic pollutant emerging from industrial and agricultural practices (Suhani et al., 2021). The system composed of the spider mites *Tetranychus urticae* and *T. evansi* feeding on tomato plants (*Solanum lycopersicum*), which hyperaccumulate cadmium in their tissues, is well characterized. Previous work showed that high concentrations of cadmium accumulation resulted in decreased performance of the two spider mite species (Godinho et al., 2018, 2022). *T. urticae* relies on several environmental cues to avoid predators (Pallini et al., 1999), pathogens (Zélé et al. 2019, but see Rodrigues et al. 2022), conspecific (Pallini et al., 1997) and heterospecific competitors (Godinho, et al., 2020a). However, whether they avoid metal-accumulating tomato plants and, if so, whether this hinges upon the environmental context, remains an open question.

## Material and Methods

### Plant and mite rearing conditions

Tomato plants (*Solanum lycopersicum*, *var* MoneyMaker) were sown in a climatic room (25 °C, 61% of humidity, light:dark = 16:8), in pots with soil (pH 5.0-6.0; SIRO, Portugal) and watered twice a week with 100 mL of tap water for the two first weeks. Subsequently, plants were transplanted to a mixture of 1:4 vermiculite, gardening soil (SIRO, Portugal), where they grew for three more weeks and were watered twice a week with 100 mL of either distilled water or a cadmium solution (3 mM or 2 mM according to the experiment, see below). All plants were also watered once a week with tap water to avoid the lack of nutrients. Just before the installation of the set-up, each plant was watered with 70 mL of tap water. This concentration of cadmium added to the soil ensures that the amount accumulated by tomato plants in their leaves is higher than the hyperaccumulating threshold (Godinho et al., 2018) and falls within values encountered in natural populations of hyperaccumulating plants (Meyer et al., 2015).

The populations of spider mites used in this study were created in 2018 by performing controlled crosses between three populations of *T. urticae* and four populations of *T. evansi* collected in the field of Portugal in 2017 from tomato plants (Godinho et al., 2020b). The resulting outbred populations have been maintained in high numbers (> 1000 adult females) in cages (28 x 39 x 28 cm) each containing two entire cadmium-free tomato plants, under controlled conditions (25 °C ± 1, 68% of humidity, light: dark = 16:8). For the experiment on individual choice and performance, populations were maintained in 5 replicates during 14 discrete generations with 220 adult mites being transferred every 14 days, as part of a larger experiment. To ensure that females used in the experiments were approximately of the same age, adult females where isolated on separate tomato leaves in plastic boxes (14 x 20 x 14 cm) and allowed to lay eggs for 48 hours. In all experiments, we used adult females resulting from this cohort, 14 days after egg laying.

The animals used in this research are not vertebrates or invertebrates with ethical and care limitations.

### Experimental set-up for individual choice and performance

To test whether spider mites individually choose between leaf discs with or without cadmium, mated females were placed at the centre of plastic strips (1×3 cm) connecting two leaf discs (Ø 16 mm) placed on soaked cotton wool, one made from a tomato plant supplied with cadmium (2 mM) and the other from a plant without cadmium (Figure 1A). For each replicate, leaf discs were made from leaves of the same age (either the 3^rd^ or 4^th^ counting from the cotyledons) and the age and position of the disc with cadmium was randomized across trials. Females (N = 17 to 24 per replicate population; total N = 101) were allowed to choose between leaves with or without cadmium for 24 hours, after which the position of the female was registered. To measure the effects of each environment on spider mite performance, females (N = 94 for plants with no cadmium and N= 97 for plants with cadmium) were isolated on leaf discs (Ø 18 mm) made from leaves 3 to 5 (from the cotyledons), from plants supplied with 2 mM cadmium chloride or not. These leaf discs were kept on top of soaked cotton wool in petri dishes. 48 hours later, females were sacrificed, and the number of eggs was counted. Assays were divided in blocks consisting of four consecutive days.

**Figure 1.**
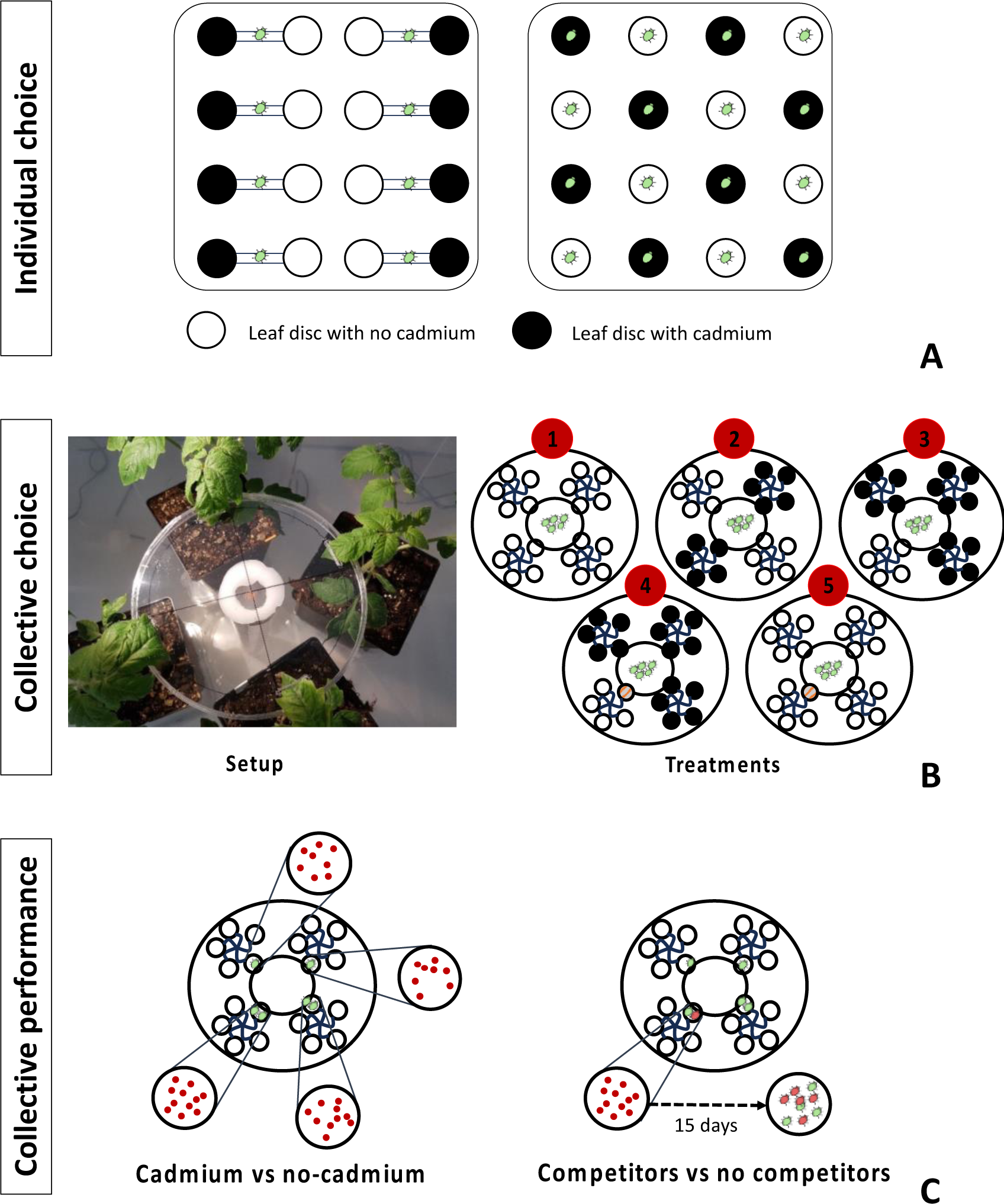
Experimental set-ups to measure individual (A) and collective host choice (B) and performance (C). The white and black shapes represent plants without and with cadmium, respectively. A) Experimental set-up for individual host choice (left panel) and individual performance (right panel). B) Experimental set-up for collective host choice and performance and schematic representation of the 5 treatments. The leaf marked in red streaks (treatments 4 and 5) identifies the presence of competitors. C) Experimental set-up for the collective performance. Depicted are treatments 1 (left panel) and 5 (right panel), which respectively represent the controls for the cadmium vs no-cadmium test and the competitors vs no competitors test, respectively. In the former, performance was assessed by directly counting the number of eggs laid on each plant. For the competitors vs no competitors test, the number of female offspring of each species was scored after 15 days, because eggs from both species are not distinguishable.

### Experimental set-up for collective choice and performance

To measure how the presence of competitors and the frequency of plants with cadmium (3 mM) affected the collective choice of spider mites, four entire tomato plants were placed in a plastic box and they were connected by a round plastic arena (15 cm of diameter) in which their 4^th^ leaf was inserted in a hole of 2 cm long at equal distance from each other and from the centre of the arena (Fig. 1B). We applied a ring of Vaseline around the petiole of the 4^th^ leaf close to the node to prevent mite dispersal to other leaves. We then released 200 females of *T. urticae* in the centre of the plastic arena, from where they could move to any plant, and counted the number of mites on each plant 24 hours later. This set-up was used to test whether spider mite distribution was random or affected by the choice of conspecifics (i.e., if there was aggregation), by giving mites the choice between four uninfested cadmium-free plants (treatment 1; Figure 1B). Additionally, we used this set-up to address how the distribution of spider mites was affected by the frequency of contaminated plants and the presence of a competitor in contaminated and uncontaminated environments. To address the first question, we placed mites in set-ups with either two (treatment 2) or three (treatment 3) plants with cadmium (Figure 1B), with the remaining plants being cadmium-free. To answer the second question, we created two other treatments, in which the leaf of one plant without cadmium was infested with 100 adult females of the competitor species *T. evansi* (50 females on the first leaflet and 25 on the two lateral leaflets to avoid overexploitation by the competitor). This plant was surrounded by three plants either with (treatment 4) or without cadmium (treatment 5; Fig. 1B). The 200 *T. urticae* females were placed in the centre of the plastic arena 24 hours after the installation of *T. evansi*. The number of *T. urticae* females on each plant was counted 24 hours later.

To measure the performance of *T. urticae* on plants with or without cadmium, the number of eggs on each leaf was counted on all plants in treatments 1, 2 and 3 after having counted (and removed) the females to measure choice. The number of eggs was then divided by the number of females on the plants, to obtain the per capita oviposition rate. To measure the performance of *T. urticae* in the presence of competitors (treatments 4 and 5), we quantified the number of adult female offspring after fifteen days, because both the eggs and the males of the two species are not distinguishable. To this aim, we removed the females from the infested leaf of the plant with competitors (from treatments 4 and 5) or from one plant without cadmium in the respective controls (treatments with the same proportion of plants with cadmium, 3 and 1, respectively – in the latter case the plant was randomly selected). Each leaf was detached and placed in a pot with water in an isolated box. Fifteen days later, the number of adult females was counted. We only report the number of *T. urticae* females. Twenty replicates per condition were done (except for treatment 5 for which 19 replicates were done).

### Statistical analysis

#### Individual choice and performance

To test if individual females choose between cadmium or cadmium-free leaf discs, a general linear mixed model with a Binomial error distribution was used. The dependent variable was the presence or absence (coded as 1 or 0, respectively) of the female in each environment (using the cbind function in R). To test the effect of cadmium on female performance, the number of eggs laid on each patch was analysed using a general linear model with a Poisson error distribution. The dependent variable was the number of eggs per patch and the test environment (cadmium-free vs. cadmium leaf discs) was coded as a fixed factor. Leaf age and replicate were used as random factors.

#### Collective choice and performance

We first tested whether spider mites distribute themselves evenly across cadmium-free plants. To this aim, we performed a G-test goodness of fit per replicate, assuming an even distribution of mites across plants (*i.e*., 25% per plant). To control for multiple tests, a Bonferroni correction was applied. Moreover, we performed a general linear model comparing the control (treatment 1) to a mock treatment where each plant ‘received’ 50 individuals of the 200 released.

All statistical analyses to test the collective spider mite behaviour were done using a generalized linear model with number of individuals on a plant as the dependent variable. For each replicate, the distribution of mites among plants was estimated in relation to the total number of mites recovered on all plants in that replicate. All models included block and the interaction between plant identity and replicate as random factors. Models where initially fit with a Poisson distribution, but due to overdispersion, a negative binomial error distribution was used instead.

To test whether spider mites collectively avoided plants with cadmium, we compared the distribution of females among plants with and without cadmium, in treatment 2, 3 and 4, using a model with cadmium and treatment and their interaction as fixed factors. Post-hoc contrasts were performed to test whether spider mites avoided plants with cadmium in each treatment, and to test whether treatments differed concerning this avoidance. To test if the frequency of plants with cadmium affected spider mite choice, we compared treatments 2 and 3, and to test if presence of competitors affected the choice of plants with cadmium we compared treatments 3 and 4. Finally, we tested whether the presence of cadmium in the environment affected the avoidance of competitors by comparing treatments 4 and 5 with a model with competitors and treatment as fixed factors and their interaction. Post-hoc contrasts were performed to test whether spider mites avoided plants with competitors in each treatment, and whether treatments differed concerning this avoidance.

All statistical analyses to test collective spider mite performance were done using general linear models with a gamma error distribution, and the number of per capita eggs or female offspring (for the treatments with competitors) as dependent variables. All models included block and replicate as random factors.

We first assessed spider mite performance in the control treatment (i.e., treatment 1) to test whether mite oviposition rate was even across plants, using a generalized mixed model with plant as fixed factor. The effect of cadmium on spider mite performance was assessed by comparing the per capita oviposition rate across all plants among treatments containing plants with and without cadmium (i.e., treatments 2, 3 and 4). For that we applied a model with cadmium and treatment and their interaction as fixed factors. Next, we performed pairwise comparisons to test whether spider mite performance differed on plants with or without cadmium within each treatment. We also performed pairwise comparisons to test whether the performance on plants with cadmium differed across treatments, only using data from plants with cadmium in each treatment.

To estimate if the effect of cadmium was similar for the individual and collective tests, we calculated the standardized effect size for each treatment and experiment using the effectsize package (Ben-Shachar et al., 2020).

To test the effect of the competitor on the number of per capita offspring we used a model with presence (or absence) of competitors as fixed factor. We used pairwise post-hoc tests with an fdr correction, to compare the treatments with competitors and their respective control (i.e. treatments 1 vs 5 and 3 vs 4).

All statistical analyses were performed using the software R v3.6 (R Core Team, 2022), using packages glmmTMB and lme4 to perform the general linear models (Bates et al., 2015; Brooks et al., 2017), car for the ANOVA tests (Fox & Weisberg, 2019), emmeans for post-hoc contrasts (Lenth, 2023) and ggplot2 for visualization (Wickham, 2009).

## Results

### Individual choice and performance

Individual spider mite females did not occupy differently leaf discs with and without cadmium (estimated overall probability = 0.0554, z ratio= 1.064, P = 0.2875, Fig. 2A). They laid significantly fewer eggs on leaf discs with cadmium than on leaf discs without cadmium (ratio between water and cadmium= 2.1, z ratio= 10.432, p<0.001, Fig. 2B, Table S1).

**Figure 2.**
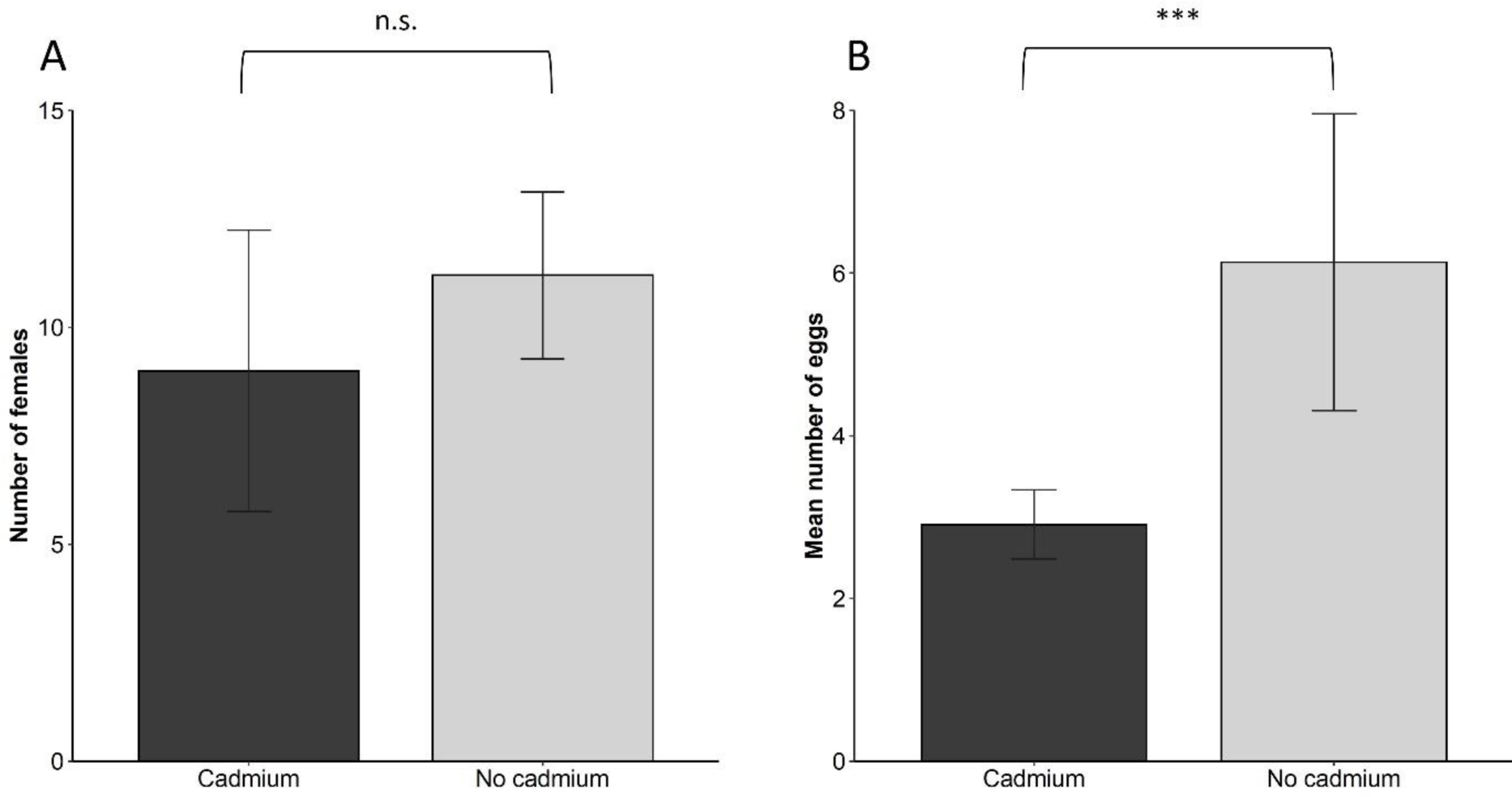
Individual host choice and performance. (A) Number of females on leaf discs from plants with cadmium (dark grey) or without cadmium (light grey) after 24 hours. (B) Number of eggs laid on leaf discs from plants with or without cadmium. Error bars represent standard error among populations (N= 17 to 24 per population; 5 populations per test). *** indicate significant differences and n.s. non-significant differences.

### Collective choice and performance

Spider mites distributed themselves unevenly across plants. In 18 out of the 19 replicates, they showed a distribution significantly different from the expected 25%, after Bonferroni correction (Table S2). Indeed, the difference between the treatment with only cadmium-free plants and no competitors and a mock variable with an even distribution across plants was significant (effect of treatment: X^2^ =7410,5; P < 0.0001, Fig. 3, treatment 1).

**Figure 3.**
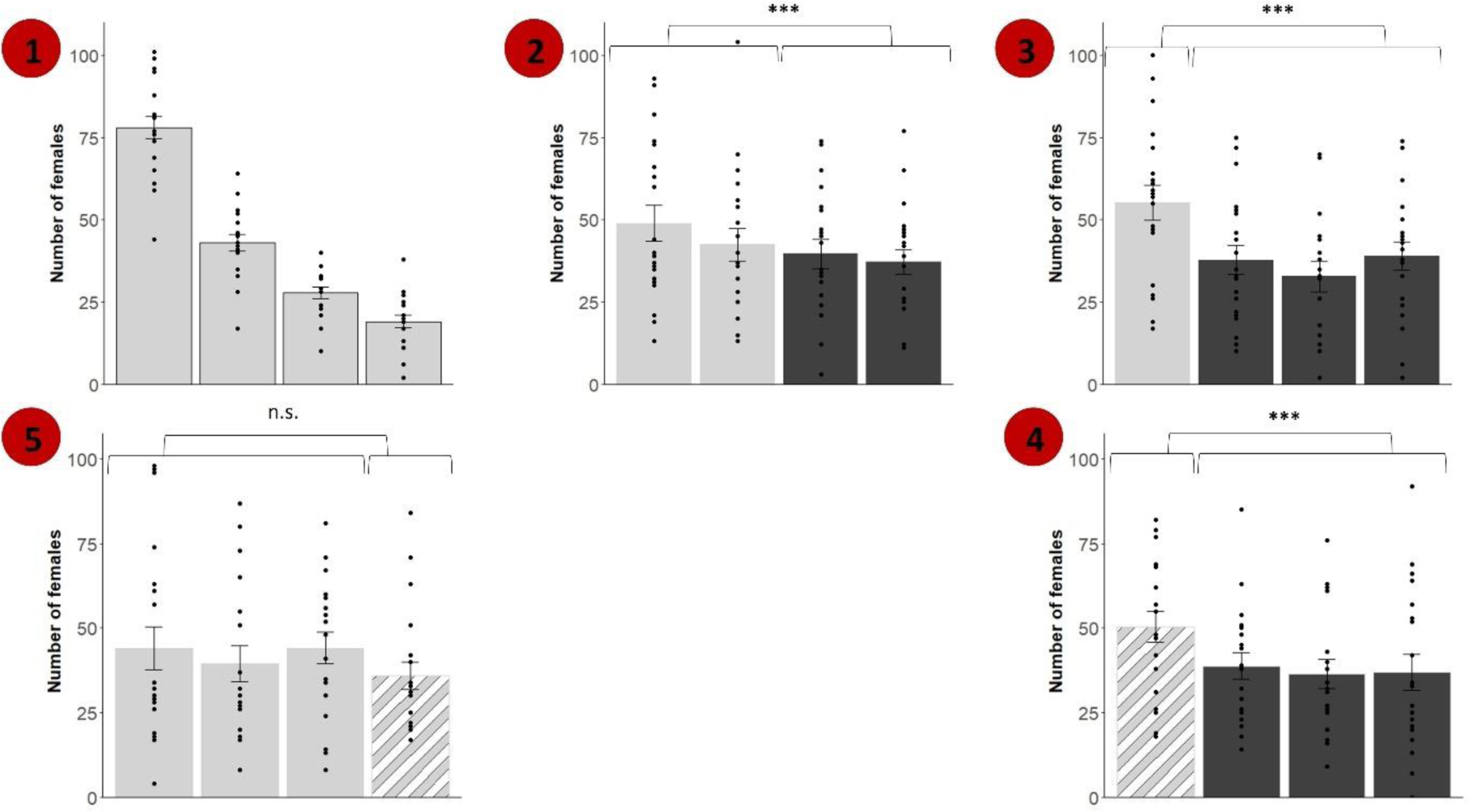
Collective host choice. Number of females after 24h on each plant for all treatments. Numbers indicate the treatments: 1) control, 2) two plants with cadmium, 3) three plants with cadmium, 4) one plant infested with competitors and three plants with cadmium and 5) one plant infested with competitors and all plants without cadmium. In treatment 1, for each replicate, plants were ranked according to the number of mites present. Depicted is the average number of mites on plants from ranks 1 to 4 (from left to right). Dark grey bars represent plants with cadmium, light grey colour plants without cadmium, white stripes represent presence of competitors on the plants. Error bars represent standard error between replicates (N= 19). *** indicate significant differences and n.s. non-significant differences.

The presence of cadmium significantly affected the number of females on each plant, but this did not differ among treatment (ANOVA results cadmium: X^2^ =8.70, p-value =0.0031; treatment: X^2^ = 1.678, p-value= 0.432; cadmium*treatment: X^2^ = 1.674, p-value = 0.432; Fig. 3). Spider mites avoided plants with cadmium in all treatments (treatment 2 plants: z ratio=-55.987, p-value <0.0001, treatment 3: z ratio=-49.560, p-value <0.0001; treatment 4: z ratio=-49.343, p-value <0.0001). This avoidance was not different between treatments with two or with three plants (contrast: z ratio=-0.390, p value = 0.9196), i.e. mites avoided plants with cadmium even when cadmium-free plants were rare in the environment.

The presence of a competitor, and its interaction with treatment significantly affected the number of females on each plant (competitor: X^2^ = 4.556, p-value= 0.033; treatment: X^2^ = 1.666, p-value=0.197; cadmium*treatment: X^2^ = 5.556, p-value =0.0184; Fig. 3). The number of mites on plants with or without competitors when all plants were cadmium-free did not differ significantly (contrast: z ratio=1.203, p-value = 0.229). However, in the treatment where plants without competitors had cadmium, females chose more often the plant with the competitor (i.e. they avoided cadmium), resulting in a significant difference among these treatments (contrast: z ratio= −2.134, p-value = 0.033). This choice was not different from that when the uncontaminated plant did not have competitors (contrast between treatment 3 and 4: z ratio=-0.457, p value= 0.891).

In the control treatment, spider mite oviposition rate was not significantly different among plants (X^2^_3_ = 3.473, p-value = 0.324; Figure 4). The presence of cadmium in plants, in treatments 2, 3 and 4 significantly reduced the oviposition rate of females (X^2^_2_ = 89.745, p-value <0.0001; Figure 4). However, this reduction in performance on cadmium-free plants was similar independently of the number of plants with cadmium in the environment (X^2^_2_ = 0.413, p-value = 0.813; Figure 4). In fact, the standardized effect size associated with the presence of cadmium was remarkably similar for treatments 2 and 3 (Table S1). The effect size of cadmium was larger in the individual than in the collective performance experiments but showed a clear overlap of the confidence intervals (Table S1).

**Figure 4.**
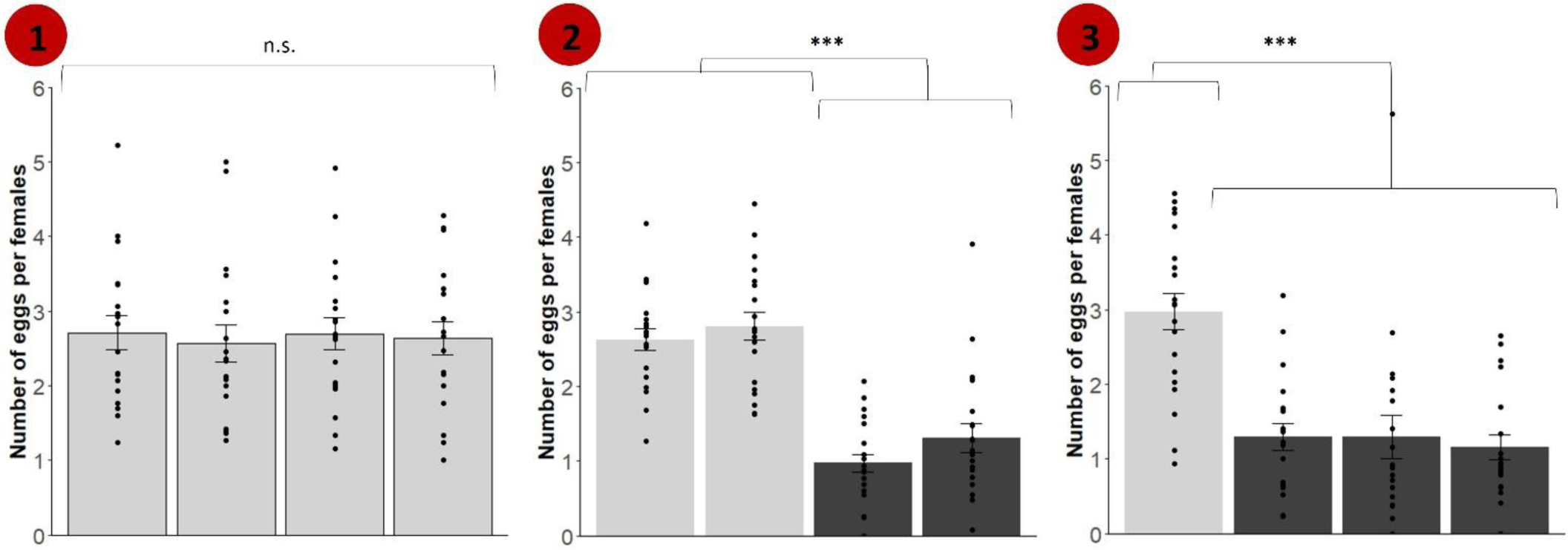
Collective performance. A) Number of eggs per female after 24h on each plant for treatments 1-3. Dark grey bars represent plants with cadmium, light grey colour plants without cadmium. Error bars represent replicate standard error between replicates (N= 19). *** indicate significant differences and n.s. non-significant differences.

In environments with cadmium-free plants only, the presence of the competitor significantly affected the number of offspring per female after one generation (X^2^_1_ = 15.877, p-value = 0.0004, Figure 5). Contrasts between plants with or without competitors revealed that the number of female offspring on uninfested plants (*i.e*., in the absence of competitors or cadmium) was higher than on cadmium-free plants with competitors (contrasts between treatment 1 and 5: t ratio= 3.771, p-value=0.002). However, this difference was not observed when competitor-free plants contained cadmium (contrasts between treatment 3 and 4: t ratio= −1.713, p-value=0.325).

**Figure 5.**
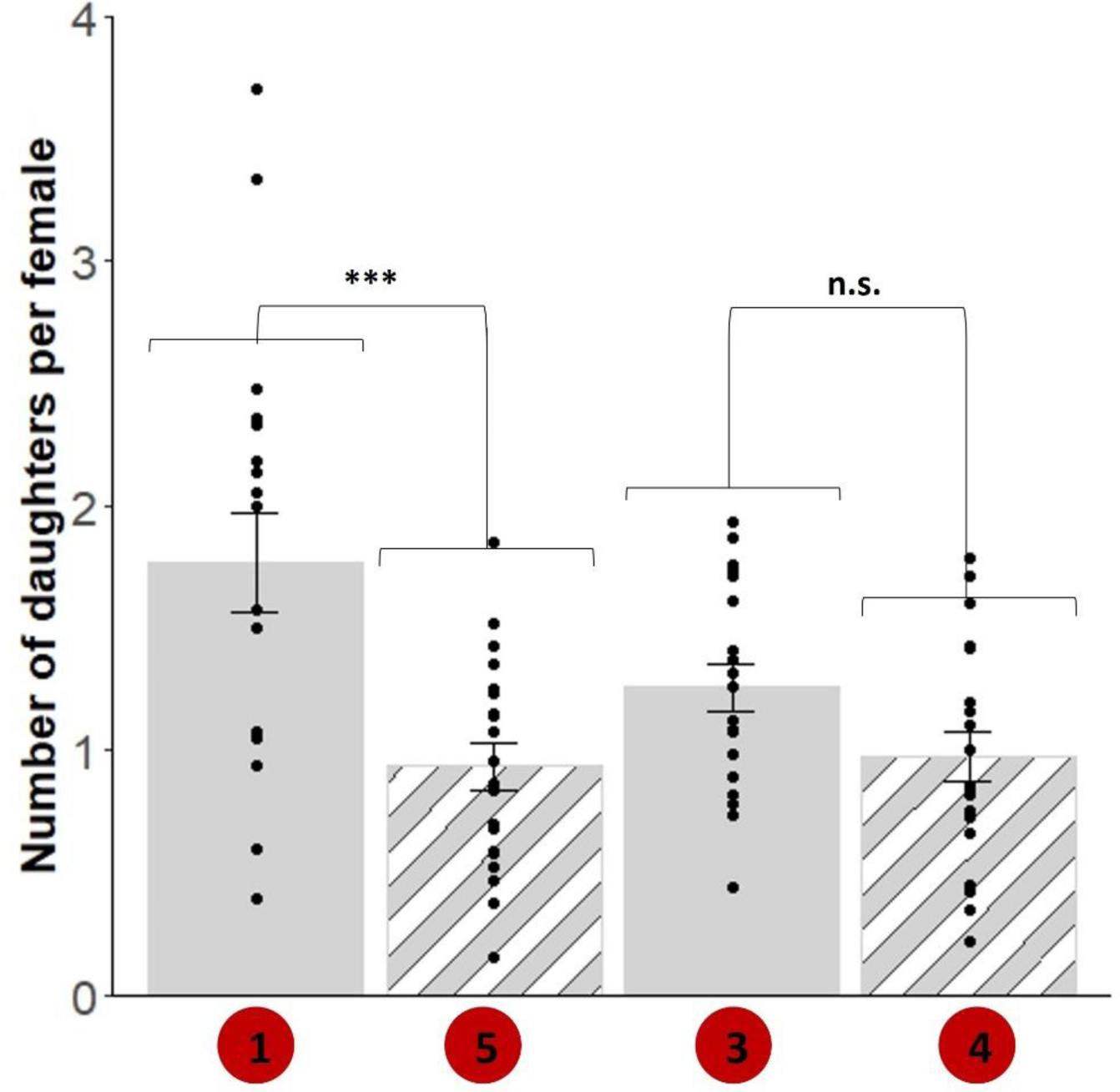
Collective performance in the presence of competitors. Given are the number of daughters per female on the plant with competitors (treatments 4 and 5) and on a plant with no cadmium of the treatments with no competitors but with the same proportion of plants with cadmium as in these treatments (i.e., no plant with cadmium: treatment 1 vs 5 – in this case the plant was randomly selected; 50% of plants with cadmium: treatment 3 vs 4). Light grey bars represent plants without cadmium and white stripes represent the presence of competitors on the plants. Error bars represent standard error among replicates (N= 19). *** indicate significant differences and n.s. non-significant differences.

## Discussion

Our study shows that *T. urticae* females do not significantly avoid tomato leaves with cadmium when they are given the choice individually, although feeding on these leaves entails a reduction in their performance. In contrast, when in group, facing the choice between cadmium and cadmium-free plants, females moved more often to plants without cadmium. They also showed a non-even distribution, even when all plants were uninfested and uncontaminated, suggesting aggregation behaviour. Moreover, the frequency of cadmium-free plants had no effect on spider mite avoidance. Finally, when all plants were cadmium-free, spider mites did not discriminate between plants with or without competitors, but they avoided plants with cadmium when the alternative was moving to a uncontaminated plant with competitors.

We found here that spider mites feeding on tomato plants that have hyperaccumulated cadmium have a lower oviposition rate than mites feeding on non-contaminated plants. This recapitulates previous findings (Godinho et al., 2018, 2022, 2024), and implies that they are robust to the set-up used. Curiously, however, *T. urticae* females did not choose between leaf discs with or without cadmium, a result also previously obtained for *T. evansi* (Godinho et al., 2024). This is surprising, not only because of the abovementioned consequences of cadmium for performance, but also because this choice set-up had been validated previously (Rodrigues et al., 2017; Zélé et al., 2019). Possibly, the presence of cadmium on the leaves does not alter their taste or volatile composition, which would inform spider mites of their content. Alternatively, such cadmium induced cues may be produced in whole plants but not in the leaf discs cut from those plants.

In contrast to the individual choice assays, spider mites systematically avoided plants with cadmium in a set-up that simultaneously tested 200 mites. Our results from the control treatment unequivocally show that spider mites aggregate on particular plants, a result also found in the field (Rijal et al., 2016). Aggregation behaviour has been shown to be beneficial to *T. urticae*, by increasing fecundity and decreasing the death rate of females living in groups (Le Goff et al., 2010). In the treatments containing plants with and without cadmium, such aggregation occurred on plants without cadmium. It is thus tempting to hypothesize that spider mites use the information of their conspecifics to avoid plants with cadmium. Spider mites often communicate by means of odour cues, such as pheromones when searching for mates (Regev & Cone, 1976; Rodrigues et al., 2017), avoiding predators (Choh & Takabayashi 2010; Pallini et al. 1999), or selecting hosts for oviposition (Weerawansha et al., 2024). Possibly, spider mites that have been on plants with cadmium produce aversive chemical cues, which are used by spider mites to avoid such plants. Alternatively, whole plants that have hyper-accumulated cadmium may produce volatiles that repel spider mites, or cadmium might disrupt the production of plant volatiles induced by spider mites upon their feeding, which are used by conspecifics to find infested plants (Lin et al., 2022; Pallini et al., 1997). Aggregation on plants without cadmium occurs even when these are rare in the environment, suggesting that the cues are easily detectable by spider mites.

In environments without cadmium, *T. urticae* spider mites did not avoid plants with competitors, which is at odds with competitors negatively affecting the number of offspring produced, as found here and in other studies (Fragata et al., 2022; Sarmento, et al., 2011). Possibly, avoidance of *T. evansi* is only observed in later stages of plant infestation, when resources become scarcer. Indeed, *T. evansi* is known to downregulate plant defences, which is beneficial to *T. urticae* in the short term (Godinho et al., 2016; Sarmento, et al., 2011). Moreover, *T. urticae* is able to avoid *T. evansi* within a single plant (Godinho, et al., 2020a), which may release the selection pressure to avoid whole plants.

In contrast, spider mites avoided plants with cadmium in detriment of avoiding competitors. An earlier study had shown that spider mites prefer young over old leaves, as the former result in higher oviposition rate for them, but that this preference vanishes when this leaf is infested with competitors (Godinho, et al., 2020a). Here, instead, we show that an abiotic selection pressure seems to be more relevant to spider mite host choice than the presence of competitors. The complex interaction between responses to abiotic and biotic stressors has been documented in other systems (Murphy & Loewy, 2015; Tran et al., 2019). One possible explanation for our results is that selecting hosts with *T. evansi* might be less risky than selecting plants with cadmium because *T. urticae* could still avoid *T. evansi* once on the plant by finding a leaf free of competitors (e.g.,Godinho et al., 2020a), while cadmium is accumulated in all leaves and there is no way to avoid it within a plant.

The avoidance of cadmium even in the presence of competitors implies stronger competition on cadmium-free plants. This is in line with studies showing increased intra or interspecific competition in resource patches with higher quality in heterogeneous environments (Shen et al., 2020; Wang et al., 2012). This suggests that spider mites will be exposed to different selection pressures in heterogeneous environments: on plants with cadmium, selection will favour mites that are able to physiologically cope with the toxic effects of metal, whereas on plants with no cadmium mites will face stronger intra and interspecific competition. If these two selection pressures persist for long enough, they may lead to high population differentiation within herbivores. Corroborating this hypothesis would necessitate studying populations of the two competitors occurring on sites with both types of plants (Freeman et al., 2007).

Alternatively, spider mites may not adapt to plants with cadmium, for example due to lack of genetic diversity, in which case the observed cadmium avoidance would lead to plants with such metal suffering few attacks by herbivores in the long run, as seen in other systems (Galeas et al., 2008; Noret et al., 2007). This should select for the maintenance of metal accumulation in plants, especially because spider mites do not individually avoid tomato plants, and not all mites avoid plants, even when collectively choosing. Thus, plants with cadmium will still suffer some herbivore damage. Hence, as long as the fitness cost of up-taking metal does not surpass the fitness gain of herbivore avoidance, selection to uptake metals as a defence against herbivores will be maintained.

### Conclusions

We demonstrated that spider mites do not significantly avoid tomato plants contaminated with cadmium when alone, despite having reduced oviposition on those plants. However, when choosing collectively, they avoid cadmium contaminated plants, even when clean plants are colonized by a superior competitor. These results suggest that collective choice in herbivores contributes to the maintenance of plant elemental defences, as it entails less herbivory on those plants. Moreover, they imply that hyperaccumulation of metals by plants may lead to stronger herbivore competition on nearby non-accumulating plants, thus having broad effects on metal-polluted ecosystems.

## Acknowledgements

We thank Cátia Eira, Inês Santos and Lucie de Sousa for maintaining plant and mite populations and all the mite squad for fruitful discussions. This work was financed by an ERC (European Research Council) consolidator grant COMPCON, GA 725419 attributed to SM and by FCT (Fundação para Ciência e Tecnologia) with a Junior researcher contract (CEECIND/02616/2018) attributed to IF.

## Author contribution

SM conceived the ideas and designed the methodology, with help from MCM; DG and MCM collected the data; DG, IF and SM analysed the data and wrote the manuscript. All authors contributed critically to the drafts and gave final approval for publication.

## Conflict of interest

Authors declare no conflict of interest.

## Data availability

Data is deposited in Figshare, DOI: 10.6084/m9.figshare.24511093.

## Notes

### Competing Interest Statement

The authors have declared no competing interest.

### Summary of Updates

This is the final version recommended by PCI Ecology

