## Supplementary material for "Spider mites collectively avoid plants with cadmium irrespective of their frequency or the presence of competitors": Table S1

Table S1 - Standardized effect size and confidence intervals (CI) estimated from the general linear models for the individual and collective performance assays.

|  | **Mean effect size** | **Lower CI** | **Upper CI** |
| --- | --- | --- | --- |
| Individual choice assay | -0.31 | -0.63 | 0.01 |
| Treatment 2 | -0.58 | -0.65 | -0.5 |
| Treatment 3 | -0.57 | -0.69 | -0.45 |
