## Supplementary material for "Spider mites collectively avoid plants with cadmium irrespective of their frequency or the presence of competitors": Table S2

| Table S2 - G-tests to assess differences in spider mite distribution in clean plants. Null expectation was considered 25% for each plant. Significant p-values are highlighted in italic. | | | |
| --- | --- | --- | --- |
| **Replicate** | **P-value** | **P-value after Bonferroni correction** | **X^2^** |
| 1 | *6.902E-04* | *7.308E-04* | *17.050* |
| 2 | *1.223E-11* | *2.446E-11* | *53.825* |
| 3 | *5.154E-28* | *9.277E-27* | *130.097* |
| 4 | *2.984E-13* | *7.673E-13* | *61.378* |
| 5 | *1.238E-06* | *1.485E-06* | *30.225* |
| 6 | *1.230E-20* | *5.535E-20* | *95.821* |
| 7 | *4.332E-15* | *1.559E-14* | *69.970* |
| 8 | *1.030E-13* | *3.089E-13* | *63.540* |
| 9 | *7.157E-23* | *4.294E-22* | *106.215* |
| 10 | *7.193E-13* | *1.619E-12* | *59.590* |
| 11 | *3.125E-09* | *5.114E-09* | *42.511* |
| 12 | *7.260E-09* | *1.089E-08* | *40.786* |
| 13 | *1.020E-06* | *1.312E-06* | *30.623* |
| 14 | *5.250E-08* | *7.269E-08* | *36.729* |
| 15 | 9.280E-01 | 9.280E-01 | 0.458 |
| 16 | *7.245E-26* | *6.521E-25* | *120.127* |
| 18 | *2.116E-04* | *2.381E-04* | *19.538* |
| 19 | *2.159E-09* | *3.886E-09* | *43.268* |
